# Dynamical patterns of coexisting strategies in a hybrid discrete-continuum spatial evolutionary game model

**DOI:** 10.1101/079434

**Authors:** A.E.F. Burgess, P.G. Schofield, S.F. Hubbard, M.A.J. Chaplain, T. Lorenzi

**Author notes:** **AMS subject classification:** 35Q92, 91A80, 92B05.

## Abstract

We present a novel hybrid modelling framework that takes into account two aspects which have been largely neglected in previous models of spatial evolutionary games: random motion and chemotaxis. A stochastic individual-based model is used to describe the player dynamics, whereas the evolution of the chemoattractant is governed by a reaction-diffusion equation. The two models are coupled by deriving individual movement rules via the discretisation of a taxis-diffusion equation which describes the evolution of the local number of players. In this framework, individuals occupying the same position can engage in a two-player game, and are awarded a payoff, in terms of reproductive fitness, according to their strategy. As an example, we let individuals play the Hawk-Dove game. Numerical simulations illustrate how random motion and chemotactic response can bring about self-generated dynamical patterns that create favourable conditions for the coexistence of hawks and doves in situations in which the two strategies cannot coexist otherwise. In this sense, our work offers a new perspective of research on spatial evolutionary games, and provides a general formalism to study the dynamics of spatially-structured populations in biological and social contexts where individual motion is likely to affect natural selection of behavioural traits.

## 1. Introduction

The term ‘spatial evolutionary game theory’ was coined in the 1990s to identify the branch of mathematics that employs methods and tools from game theory to study phenotypic evolution in spatially-structured populations. In classical models of spatial evolutionary games, players are distributed over a bounded domain in the physical space. The domain is modelled by a grid consisting of discrete nodes which can either host one player or be vacant. At a given time instant, every individual plays a two-player game with its immediate neighbours, and it is awarded a payoff according to its strategy. The occupancy of each node in the grid is consequently updated to follow the strategy that scored the highest payoff in the immediate environs. This mimics an evolutionary scenario in which the fittest phenotypes are favoured by natural selection.

Since the seminal papers of Nowak and May [27, 28, 29, 30], a torrent of theoretical analyses and modelling efforts have emerged which indicate that the results from spatial evolutionary games can be strikingly different from the outcomes of the same games played in well-mixed populations. This relates to the fact that distributing individuals over physical space allows them to become spontaneously organised into spatial structures according to their strategy. Besides leading to the generation of beautiful spatial patterns, such a form of spatial organisation fosters the emergence of spatial correlation, thus paving the way for scenarios which are much different from those encountered in well-mixed populations, where every player has the same probability to interact with any other player in the population.

Most of the classical models of spatial evolutionary games are cellular automata that do not account for the explicit motion of players. However, increasing attention has been given to spatial extensions of evolutionary game models that incorporate individual movement. For instance, Dugatkin & Wilson [11] and Enquist & Leimar [13] allowed individuals to migrate between patches without spatial structure. Diffusion-based dispersal of offspring was considered in [17, 22, 23, 41]. Ferriere & Michod [15] studied an explicit diffusive process in the context of the replicator equation, and then extended their approach by including a diffusive term [16]. Stochastic cellular-automaton models in which individuals can jump to a nearest empty site were developed in [20, 36, 40]. A dynamical system of reaction-diffusion type was investigated by Durrett & Levin [12]. A conditional mobility model on a lattice was presented in [9]. Helbing & Yu [19] introduced a model of success-driven migration, where individuals move to the sites with the highest estimated payoffs. Chen *et al.* [6] explored the effects of mobility when individuals interact with neighbours within a prescribed view radius. The case of heterogenous view radii was analysed by Zhang *et al.* [45]. An aspiration-induced migration mechanism – inducing individuals to move to new sites if their payoffs are below their aspiration level – was investigated by Yang *et al.* [44] and Lin *et al.* [24], while Meloni *et al.* [26] explored the effects of mobility in a population of Prisoner’s Dilemma players.

In this paper, we present a novel hybrid discrete-continuum modelling framework that takes into account two aspects which have been largely neglected in previous models of spatial evolutionary games: random motion and chemotaxis. In this framework, the chemotactic path is defined by the concentration of a diffusive semiochemical – *i.e.*, a chemical that conveys a signal from one individual to another – whose evolution is governed by a reaction-diffusion equation, while a stochastic individual-based model is used to describe the dynamics of players. Following the modelling strategy that Schofield *et al.* [34, 35] developed from the original approach of Anderson & Chaplain [1], we couple the two models by deriving the individual movement rules from a taxis-diffusion equation which describes the evolution of the local number of players.

For illustrative purposes, in this model we let individuals play the Hawk-Dove game originally developed by Smith & Price [25] to describe certain scenarios in animal conflict. In this particular game, individuals playing the dove strategy (*i.e.*, ‘doves’) avoid all conflict, and hence they retreat at once from a given conflict if the opponent escalates. In contrast, individuals playing the hawk strategy (*i.e.*, ‘hawks’) always escalate the conflict and continue until injured or until the opponent retreats.

Spatial aspects of the Hawk-Dove game were first mentioned in passing by Nowak & May [29, 30], and then discussed in more detail by Killingback & Doebeli [21], who reached the conclusion that, in the presence of spatial structure, doves can become spontaneously organised into clusters in which the benefits of mutual proximity can outweigh losses against hawks. Also, Hauert [18] considered the Hawk-Dove game on a lattice and remarked that “*Compared to mean field calculations, spatial extension generally favours the hawk strategy. Consequently, in spatially-structured populations, we would expect to observe more frequent escalations of conflicts than predicted by mean field theory’’*. Finally, the Hawk-Dove game on various spatial networks was analysed by Tomassini *et al.* [39], who reported that the abundance of doves depends crucially upon the network structure.

In these previous papers, although the individual players are distributed over a spatial domain, there is no explicit motion as such. Here, we provide evidence through computational simulations that letting individuals diffuse through space, and move chemotactically (i.e. up semiochemical gradients), can bring about self-generated patterns which create favourable conditions for the co-existence of hawks and doves in situations in which the two strategies cannot coexist either in spatially homogeneous models or in cellular automaton models.

## 2. The model

We study the dynamics of players who move in a square domain Ω ≡ [−*ℓ*, *ℓ*] × [−*ℓ*, *ℓ*]. Individual movement is seen as the superposition of spatial diffusion and chemotactic response. The former is due to random motion, whilst the latter is guided by the gradient of a semiochemical emitted by the players themselves. At each time instant, players occupying the same position can engage in a two-player game, and are awarded a payoff according to their strategy. The accumulated payoff then determines the number of offspring which can be produced by a player.

### 2.1. Movement rules

At each time instant *t* ≥ 0, the concentration of semiochemical and the number of players at position (*x, y*) ∈ Ω are characterised by the functions *S*(*t, x, y*) ≥ 0 and *P* (*t, x, y*) ≥ 0, respectively.

The evolution of *S*(*t, x, y*) is governed by the following reaction-diffusion equation

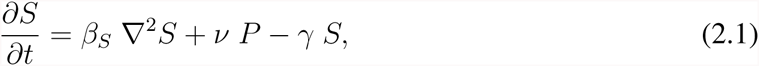

which is completed with reflective (no-flux) boundary conditions. Eq.(2.1) relies on the assumptions that the semiochemical is produced by all players at the same rate *ν* ≥ 0, undergoes a linear decay process at rate *γ* > 0, and diffuses at rate *β*_*S*_ > 0.

To describe the movement of players, we make use of the following strategy:

i. We introduce the taxis-diffusion equation below

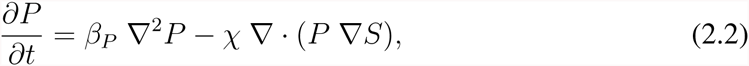

which we complete with no-flux boundary conditions. In Eq.(2.2), the diffusion term models the tendency of players to diffuse through space with motility *β*_*S*_ > 0. The advection term accounts for the fact that players move up the semiochemical gradient, and the parameter *χ* > 0 is the chemotactic sensitivity coefficient.
ii. We fix a time step ∆*t* and set *t*_*n*_ = *n*∆*t*. Also, we discretise the square Ω with a uniform mesh as

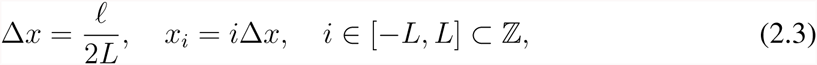

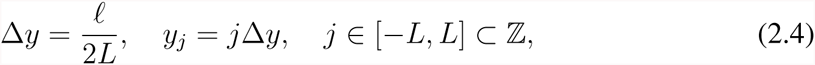

and thereafter we approximate *S*(*t*_*n*_, *x*_*i*_, *y*_*j*_) and *P* (*t*_*n*_, *x*_*i*_, *y*_*j*_) by discrete values 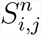 and 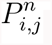, respectively.
iii. Following Schofield *et al.* [34, 35], we discretise Eq.(2.2) by using an explicit five-point central difference scheme to obtain the following algebraic equation for 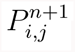, *i.e.*, the number of players at grid-point (*x*_*i*_, *y*_*j*_) at the time step *n* + 1:

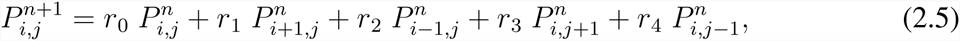

where the coefficients *r*_0,1,2,3,4_ are given by

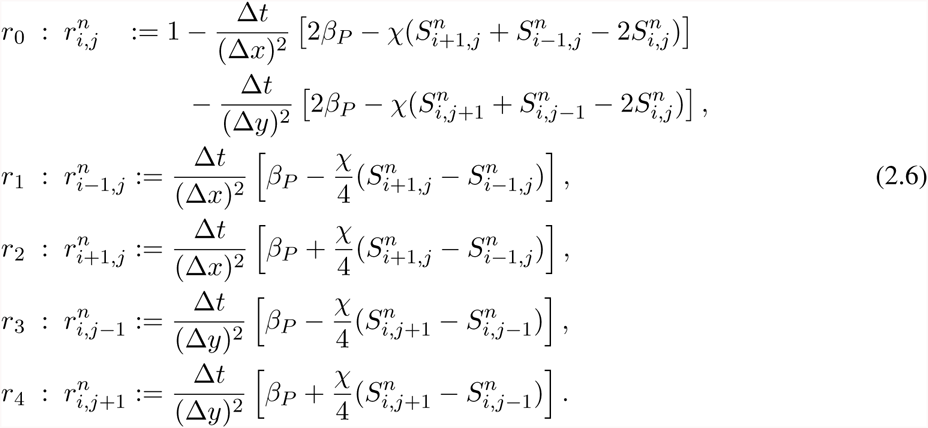

These coefficients are proportional to the probabilities of a player being stationary (*r*_0_), or moving left (*r*_1_), right (*r*_2_), down (*r*_3_) or up (*r*_4_), and hence the above system may be used to generate the movement of players from grid-point to grid-point.

In this framework, at any step *n* > 1, the algorithm for individual movement is as follows:

i. The value of 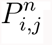 is identified by counting the number of players at every grid-point (*x*_*i*_, *y*_j_), and the semiochemical concentration is computed by calculating the numerical solutions of the mathematical problem defined by completing Eq.(2.1) with zero Neumann boundary conditions and suitable initial conditions.
ii. The probabilities 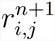 are evaluated at each grid-point by substituting the local semiochemical concentrations into Eqs.(2.6).
iii. At each grid-point (*x*_*i*_, *y*_*j*_), the values of the spatial-transition probabilities are used to define five intervals as

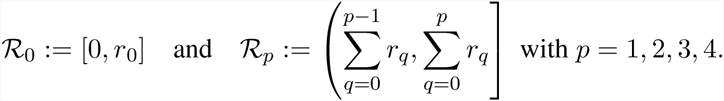
iv. For each player at a given grid-point, a random real number between 0 and 1 is generated, and a comparison of this number with the above ranges yields the direction of movement of the player. Namely, the player will not move if the random number belongs to 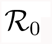, or it will move left if the number belongs to 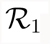, right if the number belongs to 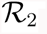, down if the number belongs to 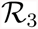 and up if the number belongs to 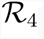.

### 2.2. Interaction and reproduction rules

At any step *n* > 1, players that occupy the same grid-point engage in a two-player game and are awarded a payoff according to their strategy. Each player can play at most *M* rounds of the game, either with the same opponent or with different opponents. We assume that all players have the same lifetime *τ*, and consider two possible underlying models for the reproduction rules:

i. a synchronous model, where players give birth a number of offspring equal to the integer part of the accumulated payoff at the end of their lives;
ii. a non-synchronous model, where reproduction occurs at each time step with an individual producing a number of offspring equal to the integer part of its current accumulated payoff, and the reproductive fitness being then decreased by this same number.

In both cases, offspring are initially located at the same site as the parent individual, and they inherit its strategy. In this setting, the constraint on the maximum number of interactions introduces an indirect limitation on the number of players, since it limits the potential maximum gain in reproductive fitness.

As a specific example, we let individuals play the Hawk-Dove game, a two-player game in which each player can adopt either one of two strategies: Hawk (H) – *i.e.*, escalate and continue until injured or until opponent retreats – or Dove (D) – *i.e.*, retreat at once if opponent escalates [25]. In this case, the outcome of the game is determined by the following payoff matrix:

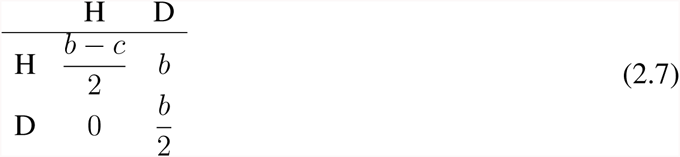

If both players play the strategy H, one wins and receives a benefit *b* > 0 with probability 1/2, while the other loses and pays a cost *c* > 0 with the same probability. Hence, the expected payoff for each player is (*b* − *c*)*/*2. If both players play the strategy D, one receives a benefit *b* > 0 with probability 1/2, whereas the other neither pays any cost nor receives any benefit. Therefore, the expected payoff is *b/*2. Finally, if one player plays H while the other plays D, the former wins and receives payoff *b*, while the latter retreats and receives payoff 0.

## 3. Simulation results

To perform numerical simulations, we choose *ℓ* = 100 and use the mesh defined by (2.3)-(2.4) with *L* = 50 (*i.e.*, ∆*x* = ∆*y* = 1) to discretise the spatial domain. Focussing on the case where a single hawk invades a population of doves, we let the initial system consist of 1 hawk located at the centre of a randomly scattered population of 4.999 × 10^3^ doves. Moreover, we assume that there is no semiochemical inside the system at *t* = 0, that is, we set

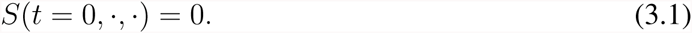

The method we use to construct numerical solutions of the mathematical problem defined by completing Eq.(2.1) with (3.1) and zero Neumann boundary conditions, is based on an explicit five-point central difference scheme for the discretisation of the diffusion term. Since ∆*x* = ∆*y* = 1, we select ∆*t* = 0.05 to meet the CFL condition, and thus ensure the stability of the numerical scheme.

We focus on a set of parameter values that is representative of an extensive range of simulation results. In analogy with the choice made by Schofield *et al.* [34], we define the semiochemical diffusion coefficient *β*_*S*_ := 5 × 10^−4^, the semiochemical decay constant *γ* := 1, the chemotactic sensitivity coefficient *χ* := 2 × 10^−4^ and the player motility *β*_*P*_ := 5 × 10^−3^. To study the dynamics of the system with or without semiochemical secretion, we alternatively use the definition *ν* := 1 or *ν* := 0. For the payoff matrix (2.7), we set *b* = 0.04 and *c* = 0.03 in the case of synchronous generations, and *b* = 0.03 and *c* = 0.04 in the case of non-synchronous generations. We set the maximum number of interactions per player per iteration *M* = 4, and we define the player lifetime *τ* := 100∆*t*. We run simulations for 5 × 10^5^ time steps.

### 3.1. Dynamics with synchronous generations

We focus here on the case in which the players proliferate in a synchronous way (as previously defined). We begin by examining the effects of semiochemical secretion on the spatial-temporal dynamics of hawks and doves. We will then investigate how the dynamics change in response to variations in the value of the parameter *b* of the payoff matrix (2.7).

#### 3.1.1. Dynamics without semiochemical secretion

In the absence of semiochemical secretion (*i.e.*, when *ν* := 0, no chemotaxis), the results in Figure 1 and Figure 2 show that the proliferation of the central hawk gives rise to an almost circular wavefront made up of hawks, whose expansion causes a rapid and drastic decline in the local density of doves. A substantial number of hawks survive in the space behind the invasive front, leading to the formation of a trailing edge of hawks. In the trailing edge, hawks are left without doves with which they can interact. As a consequence, they die out within a few generations. Such a process culminates in the formation of small and highly concentrated clusters of hawks and doves, which are confined to a small area of the spatial domain. Successively, the interaction between these clusters leads to the emergence of densely populated filamentary structures of doves which are chased by sparser filaments of hawks.

**Figure 1:**
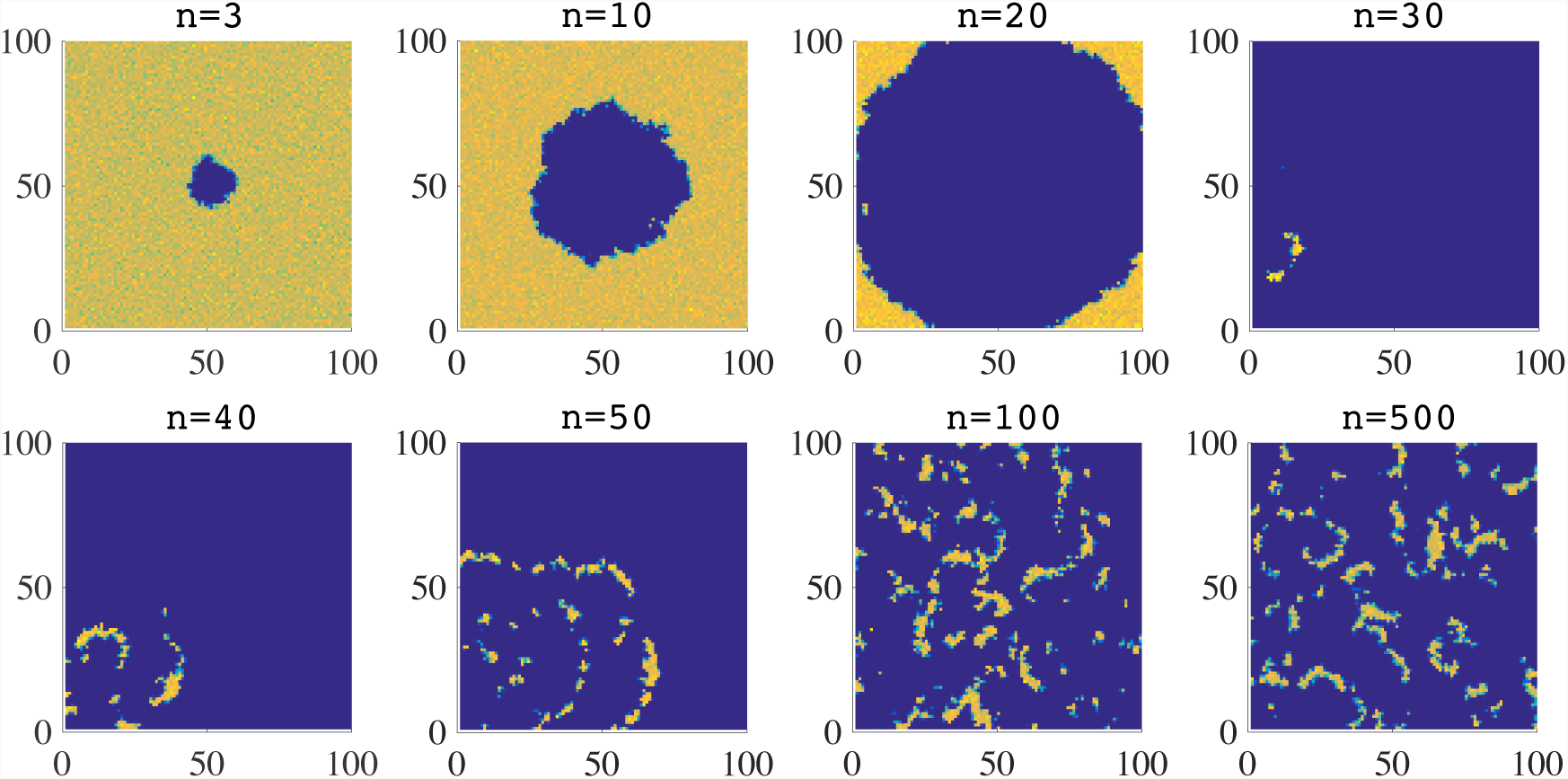
Plots summarising the time-evolution of the spatial distribution of doves, in the absence of semiochemical secretion (*i.e.*, when *ν* = 0) and with synchronous generations. The colour scale ranges from blue (low density) to yellow (high density). The time step *n* is in units of 10^3^.

**Figure 2:**
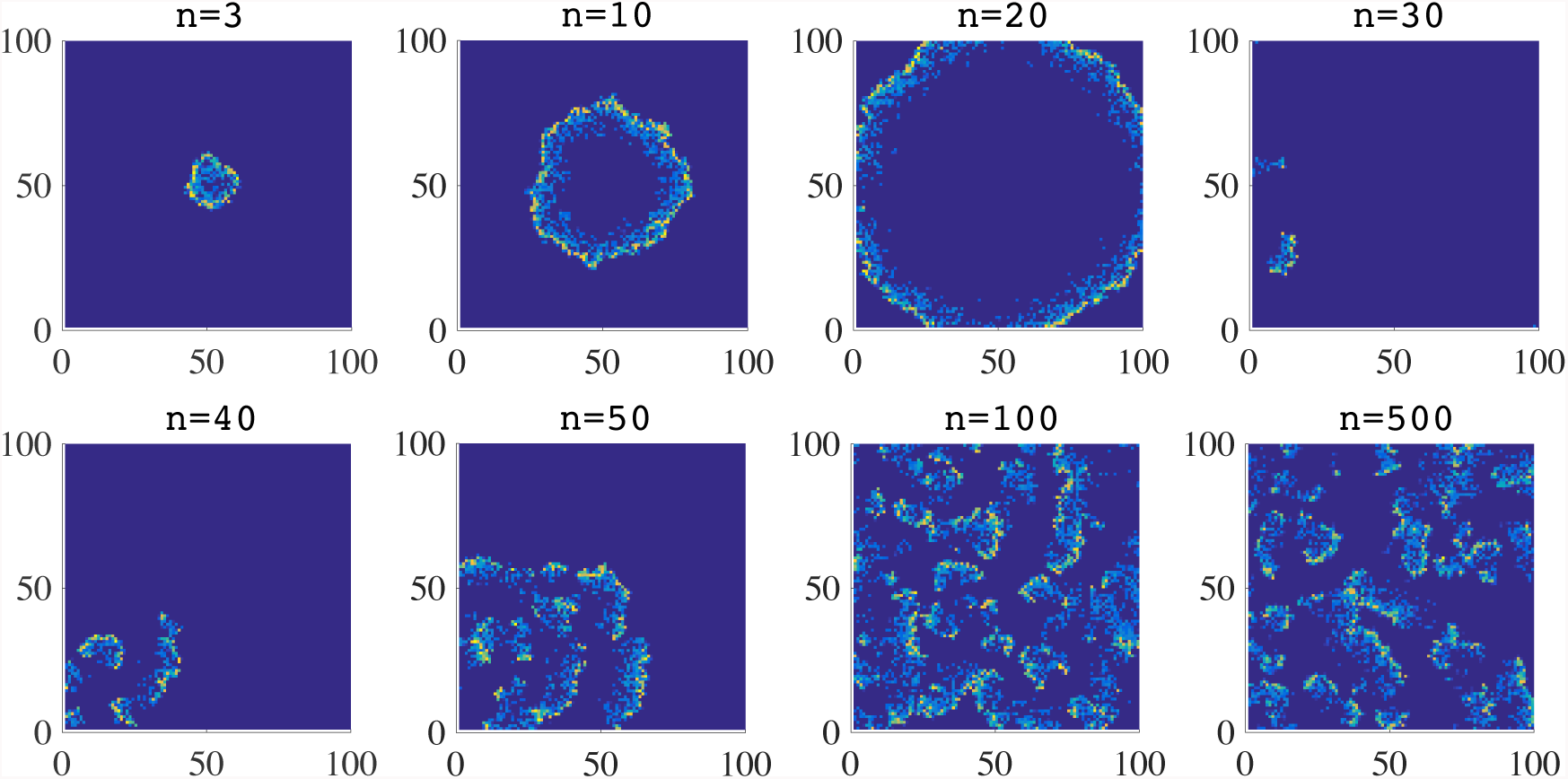
Plots summarising the time-evolution of the spatial distribution of hawks, in the absence of semiochemical secretion (*i.e.*, when *ν* = 0) and with synchronous generations. The colour scale ranges from blue (low density) to yellow (high density). The time step *n* is in units of 10^3^.

As highlighted by the results presented in Figure 3, after a severe reduction in the total number of doves at the beginning of simulations, the total numbers of doves and hawks fluctuate about well defined non-zero values, and the two strategies coexist in a stable way. The time average of the total number of doves is 3.9853 × 10^4^, while that of hawks is 1.5820 × 10^4^.

**Figure 3:**
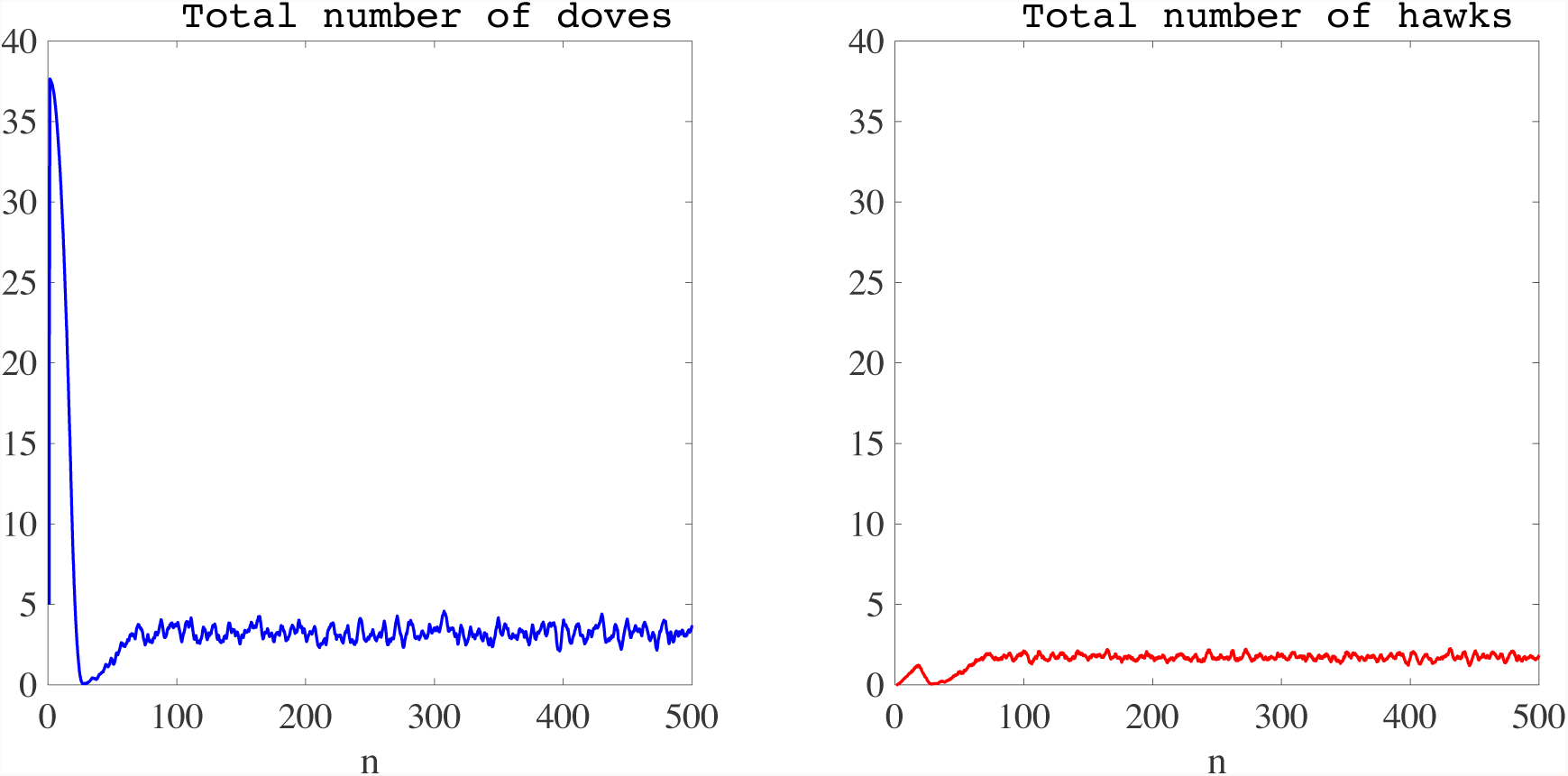
Plot of the total number of doves (left panel) and hawks (right panel) as a function of time, in the absence of semiochemical secretion (*i.e.*, when *ν* = 0) and with synchronous generations. The time average of the total number of doves is 3.9853 × 10^4^, while that of hawks is 1.5820 × 10^4^. The time step *n* is in units of 10^3^, and the total numbers of doves and hawks are in units of 10^4^.

#### 3.1.2. Dynamics with semiochemical secretion

When semiochemical is secreted by players (*i.e.*, when *ν* = 1, chemotactic movement present), in analogy with the previous case, the progeny of the single central hawk becomes progressively organised into an almost circular expanding wavefront (*vid.* Figure 4). However, compared with the case when there is no semiochemical secretion, the wavefront is thiner, and leaves in its wake a few doves that coexist with a few hawks. We observe the formation of clusters of doves which are followed closely by flocks of hawks. The interaction between doves and hawks induces a rapid decline in the local number of doves, yielding dynamical and fluctuating spatial patterns.

**Figure 4:**
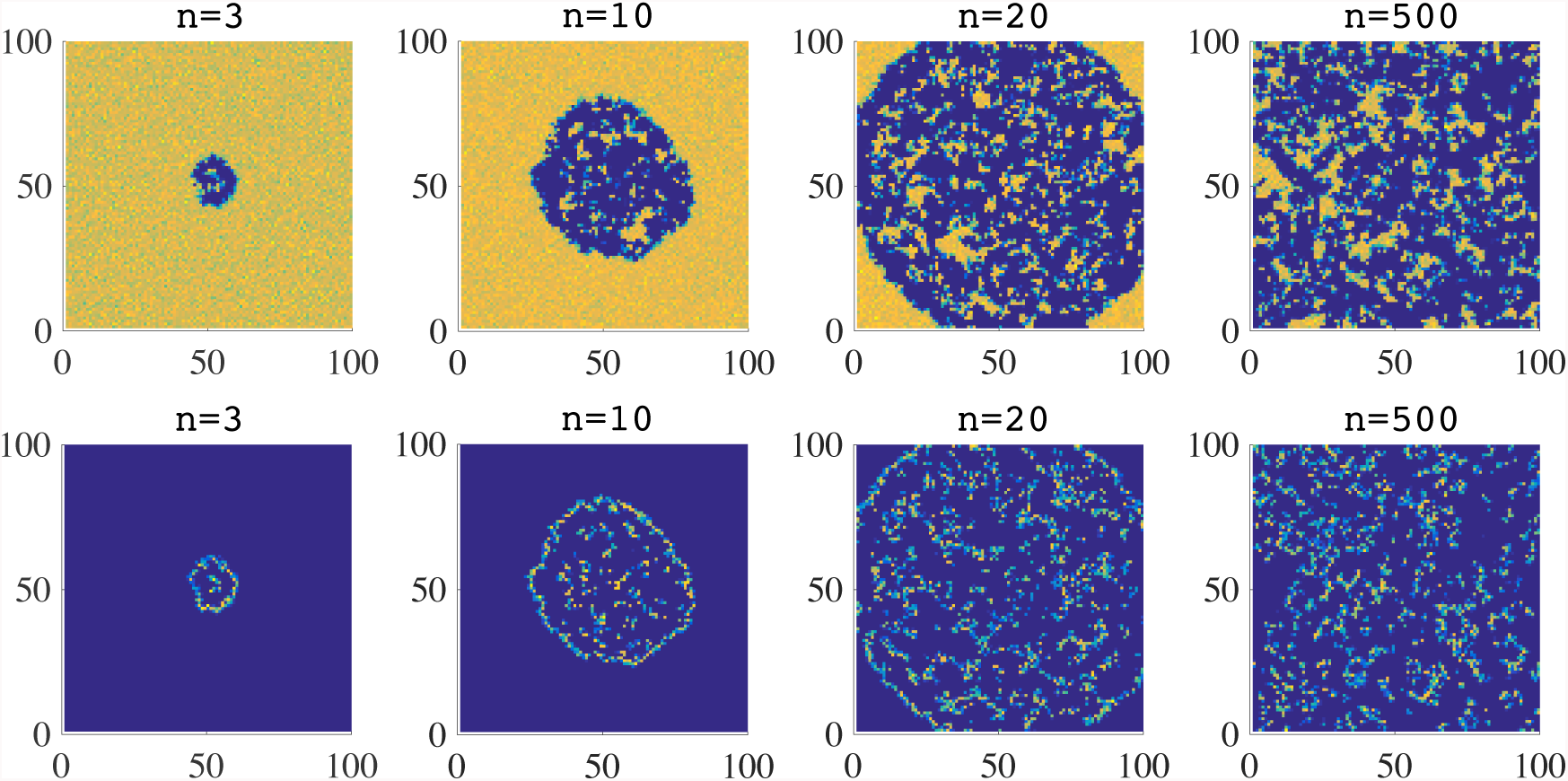
Plots summarising the time-evolution of the spatial distribution of doves (top panels) and hawks (bottom panels), in the presence of semiochemical secretion (*i.e.*, when *ν* = 1) and with synchronous generations. The colour scale ranges from blue (low density) to yellow (high density). The time step *n* is in units of 10^3^.

Once again, the strategies coexist in a stable way (see the results presented in Figure 5). The mean (with respect to time) of the total number of hawks and the equivalent mean of doves are, respectively, 10.3450 × 10^4^ and 2.1118 × 10^4^. These values are higher than those obtained in the case when there is no semiochemical secretion (compare the results in Figure 5 with the curves in Figure 3). Also, the ratio between the time average of the total number of hawks and the time average of the total number of doves is lower than that obtained without semiochemical secretion.

**Figure 5:**
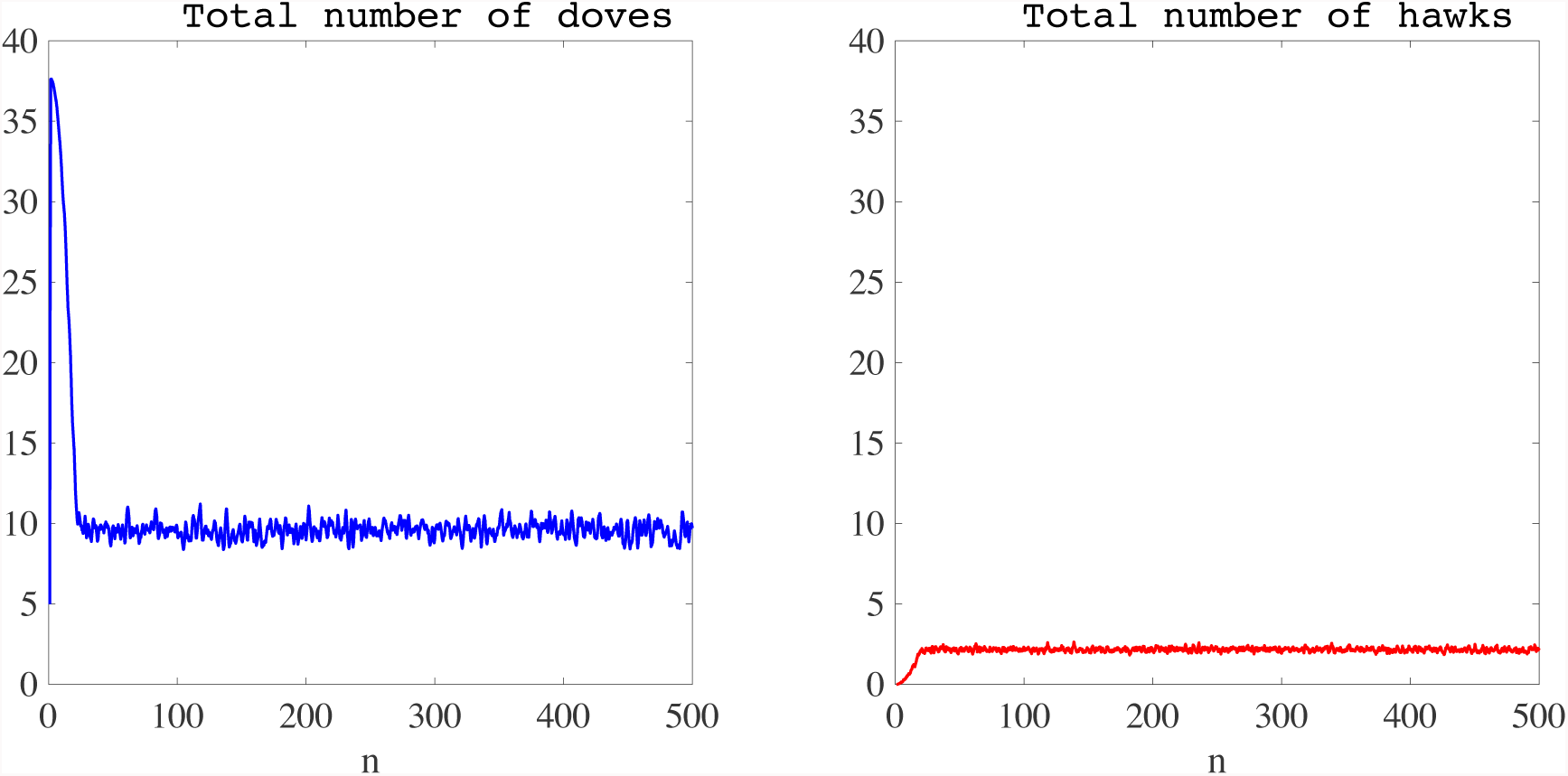
Plot of the total number of doves (left panel) and hawks (right panel) as a function of time, in the presence of semiochemical secretion (*i.e.*, when *ν* = 1) and with synchronous generations. The time average of the total number of doves is 10.3450 × 10^4^, while that of hawks is 2.1118 × 10^4^. The time step *n* is in units of 10^3^, and the total numbers of doves and hawks are in units of 10^4^.

#### 3.1.3. Effects of varying the benefit parameter *b*

Finally, to investigate how the value of the benefit parameter *b* impinges on the interaction dynamics of hawks and doves, we perform again the same simulations of the two previous subsections while holding all parameters constant except for the benefit *b*, and we record the time average of the total number of doves and hawks.

The results obtained are illustrated by the plots of Figure 6, which show how the time average of the total numbers of doves and hawks vary as a function of *b*, in the absence (left panel) or in the presence (right panel) of semiochemical secretion. For values of *b* sufficiently smaller than the cost *c*, the two strategies cannot coexist, and hawks go extinct. Again, for values of *b* sufficiently larger than *c*, all doves die out and only hawks survive. Between these extremes, we observe the stable invasion of a minority of hawks along with a substantial decrease in the average size of the resident population of doves. The limiting *b* value consistent with the stable coexistence of the two strategies is approximately the same, both in the presence and in the absence of semiochemical secretion.

**Figure 6:**
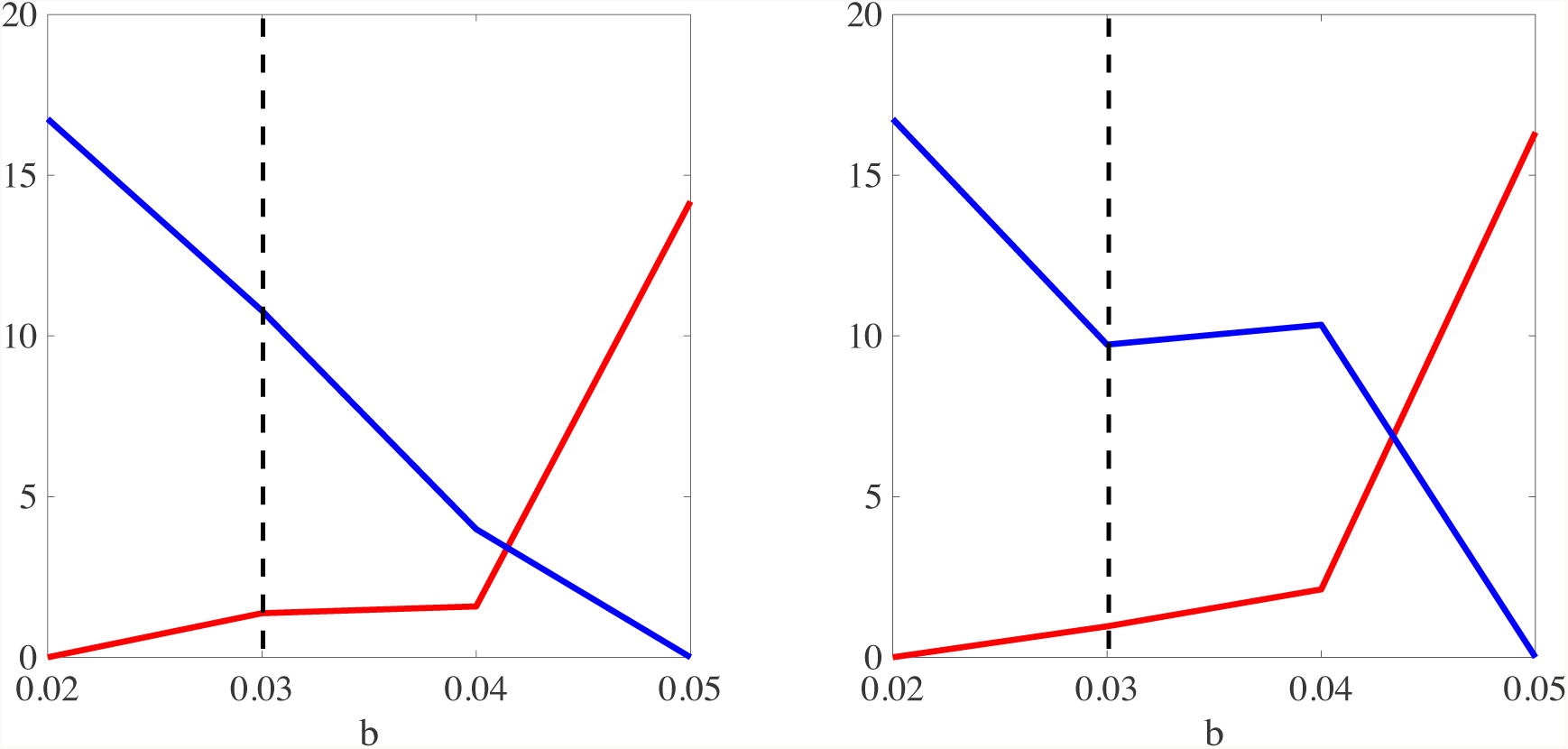
Plot of the time average of the total number of doves (blue lines) and hawks (red lines) as a function of the parameter *b*, in the absence (left panel) or in the presence (right panel) of semiochemical secretion and with synchronous generations. The black dashed lines highlight the value of the parameter *c* used to perform simulations. Time averages are in units of 10^4^.

### 3.2. Dynamics with non-synchronous generations

We now consider the scenario in which players’ generations are non-synchronous (as previously defined). In analogy with the case of synchronous generations, we study the effects of semiochemical secretion on the spatio-temporal dynamics of hawks and doves first, and then we investigate how the dynamics change in response to variations in the value of the parameter *b* of the payoff matrix (2.7).

#### 3.2.1. Dynamics without semiochemical secretion

In the absence of semiochemical secretion (*i.e.*, when *ν* := 0, no chemotaxis), the results in Figure 7 reveal that the hawks once again spread through the population of doves, but that the wave-front of invading hawks reported in the case of synchronous generations is not observed. The distribution of doves appears to be uniform over much of the grid and punctuated by numerous small empty regions of the domain. These regions emerge due to the presence of small islands of highly concentrated hawks which outcompete the resident doves.

**Figure 7:**
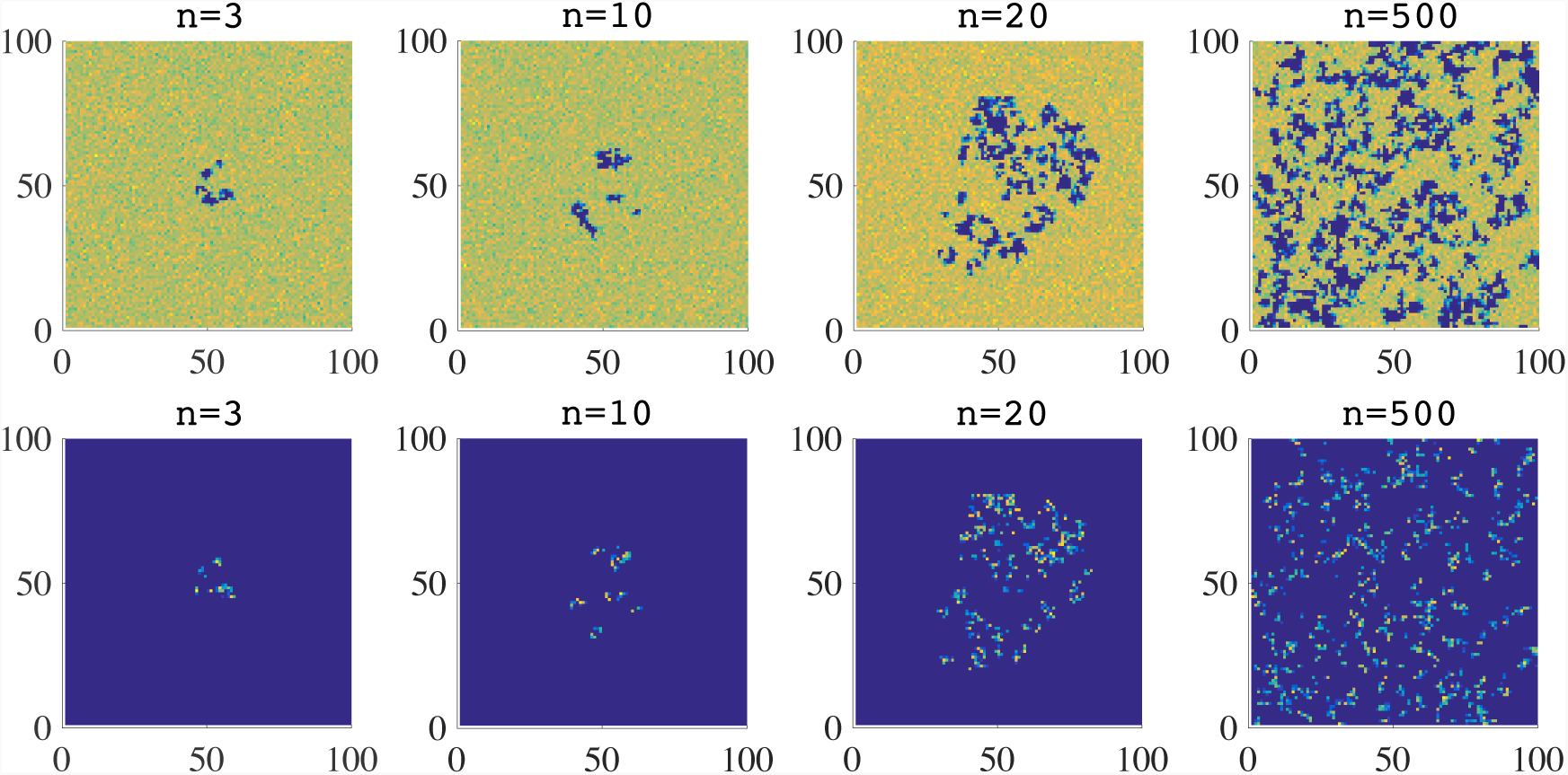
Plots summarising the time-evolution of the spatial distribution of doves (top panels) and hawks (bottom panels), in the absence of semiochemical secretion (*i.e.*, when *ν* = 0) and with non-synchronous generations. The colour scale ranges from blue (low density) to yellow (high density). The time step *n* is in units of 10^3^.

As highlighted by the results presented in Figure 8, the total numbers of doves and hawks fluctuate about well-defined non-zero values, and the two strategies coexist in a stable way. The time average of the total number of doves is 195.590 × 10^3^, while that of hawks is 7.039 × 10^3^.

**Figure 8:**
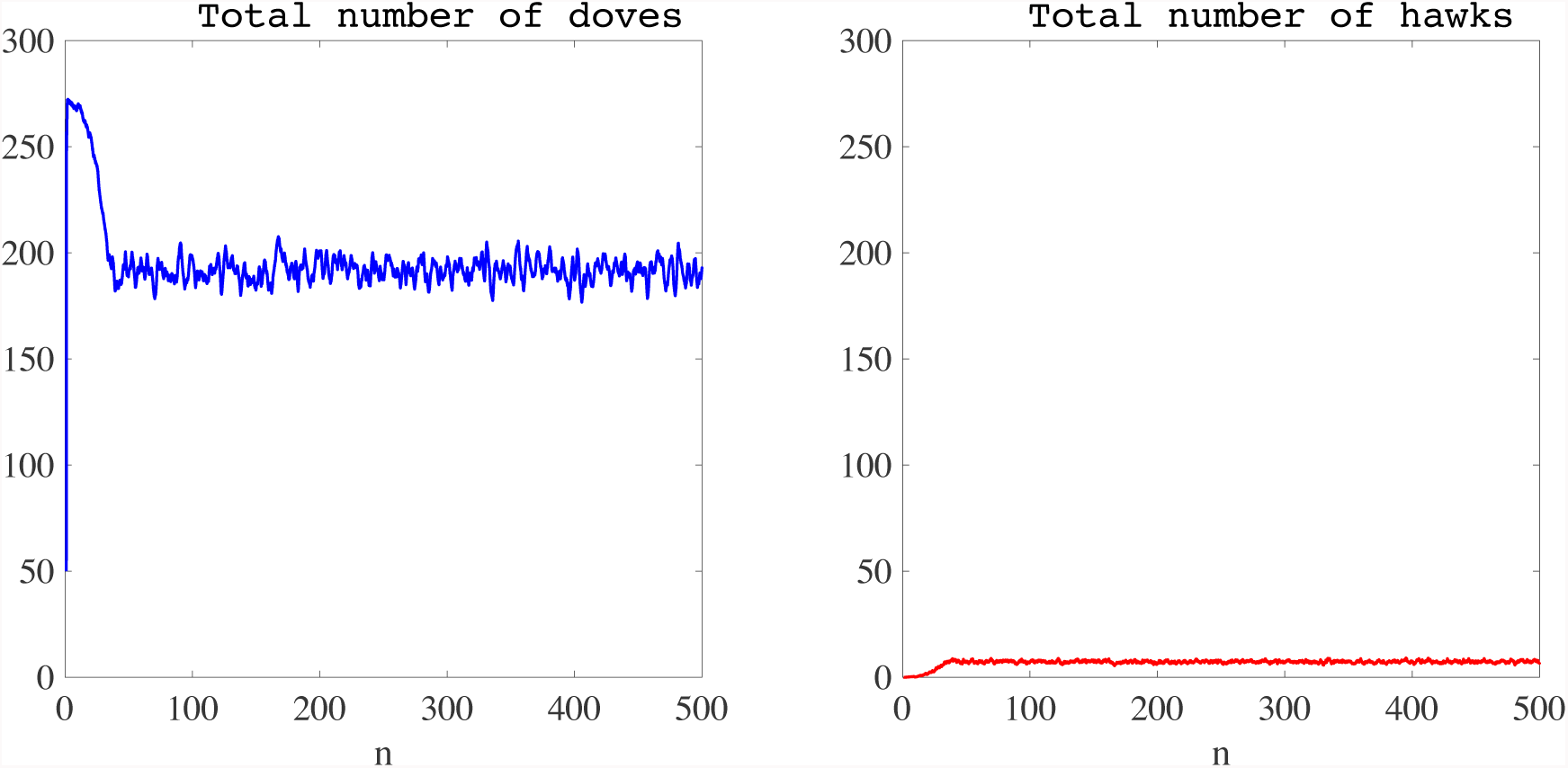
Plot of the total number of doves (left panel) and hawks (right panel) as a function of time, in the absence of semiochemical secretion (*i.e.*, when *ν* = 0) and with non-synchronous generations. The time average of the total number of doves is 195.590 × 10^3^, while that of hawks is 7.039 × 10^3^. The time step *n*is in units of 10^3^, and the total numbers of doves and hawks are in units of 10^3^.

#### 3.2.2. Dynamics with semiochemical secretion

When semiochemical is secreted by players (*i.e.*, when *ν* = 1, chemotactic movement present), the spatial dynamics of doves and hawks are similar to those observed in the case where there is no semiochemical secretion (compare the plots in Figure 9 with the plots in Figure 7).

**Figure 9:**
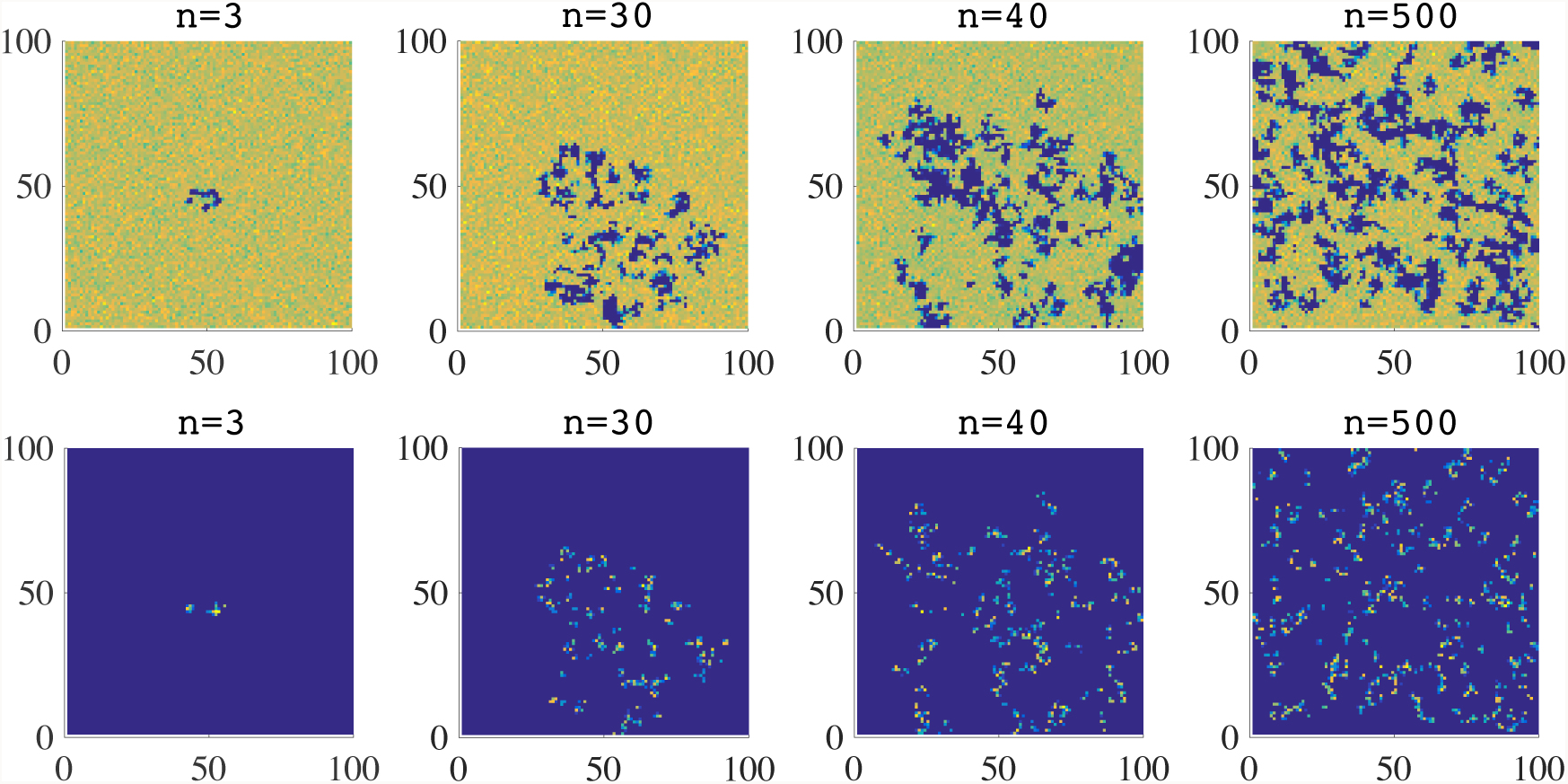
Plots summarising the time-evolution of the spatial distribution of doves (top panels) and hawks (bottom panels), in the presence of semiochemical secretion (*i.e.*, when *ν* = 1) and with non-synchronous generations. The colour scale ranges from blue (low density) to yellow (high density). The time step *n* is in units of 10^3^.

The strategies coexist in a stable way (see the results presented in Figure 10). The mean (with respect to time) of the total number of hawks and the equivalent mean of doves are, respectively, 4.631 × 10^3^ and 197.080 × 10^3^. These values are similar to those obtained in the absence of semiochemical secretion (compare the results in Figure 10 with those in Figure 8). Also, the ratio between the time average of the total number of hawks and the time average of the total number of doves is lower than that obtained without semiochemical secretion.

**Figure 10:**
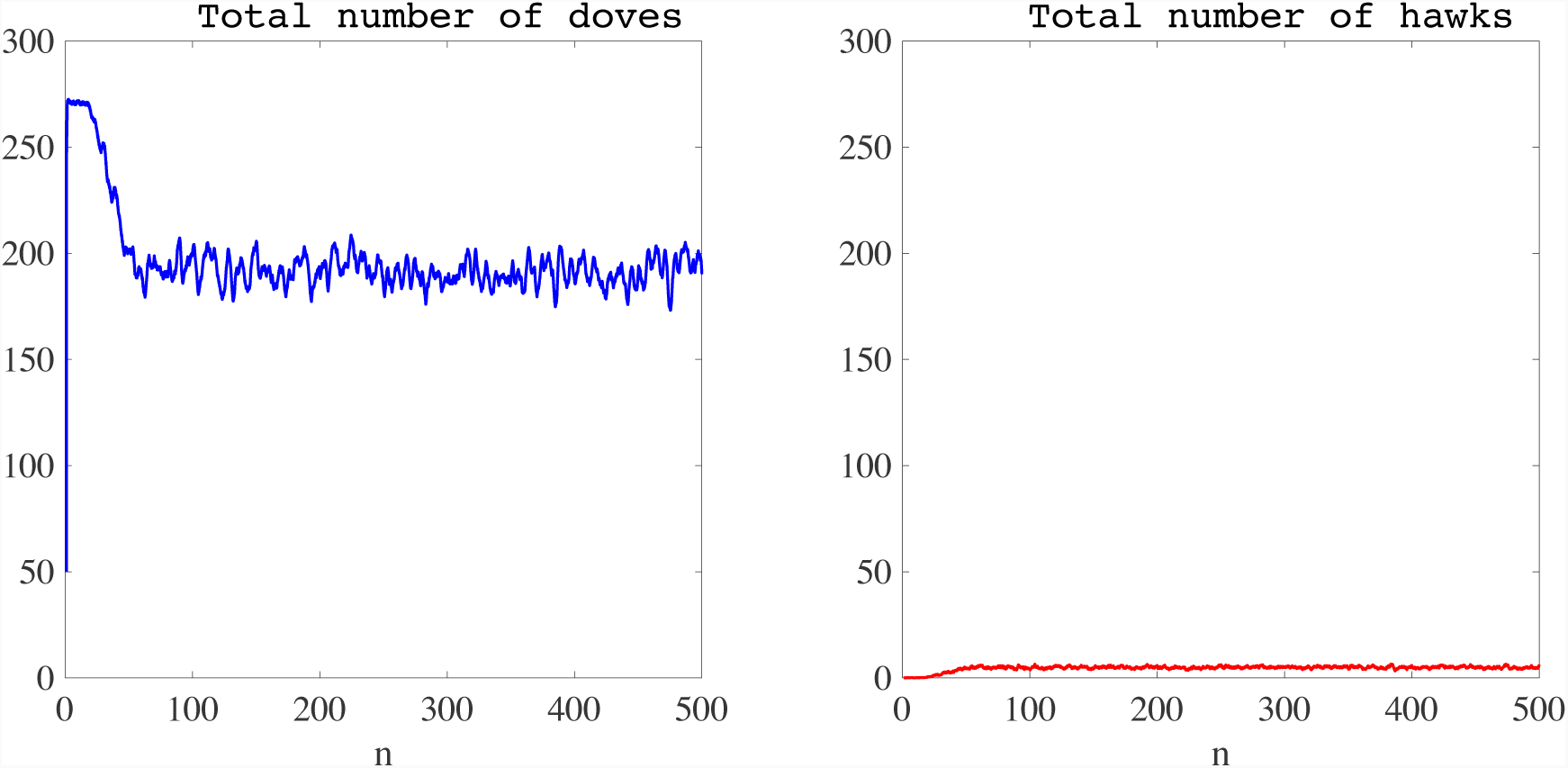
Plot of the total number of doves (left panel) and hawks (right panel) as a function of time, in the presence of semiochemical secretion (*i.e.*, when *ν* = 1) and with non-synchronous generations. The time average of the total number of doves is 197.080 × 10^3^, while that of hawks is 4.631 × 10^3^. The time step *n* is in units of 10^3^, and the total numbers of doves and hawks are in units of 10^3^.

#### 3.2.3. Effects of varying the benefit parameter *b*

As we did in the case of synchronous generations, now we perform simulations holding all parameters constant except for the benefit *b*, and we record the time average of the total number of doves and hawks. To allow for comparison with the results obtained for synchronous generations, we set *c* = 0.03. The results obtained are presented in the plots of Figure 11, which show how the time average of the total numbers of doves and hawks vary as a function of *b*, in the absence (left panel) or in the presence (right panel) of semiochemical secretion. In analogy with the case of synchronous generations, for values of *b* sufficiently smaller than the cost *c*, the two strategies cannot coexist, and hawks go extinct. Again, for values of *b* sufficiently larger than *c*, all doves die out and only hawks survive. Between these extremes, the situation is more complicated than in the case of synchronous generations. In fact, there exist intermediate values of *b* for which the stable coexistence of the two strategies occur but there are also intermediate values of *b* that bring about the co-extinction of hawks and doves. We note that the range of *b* values where co-extinction is present becomes wider in the presence of semiochemical secretion

**Figure 11:**
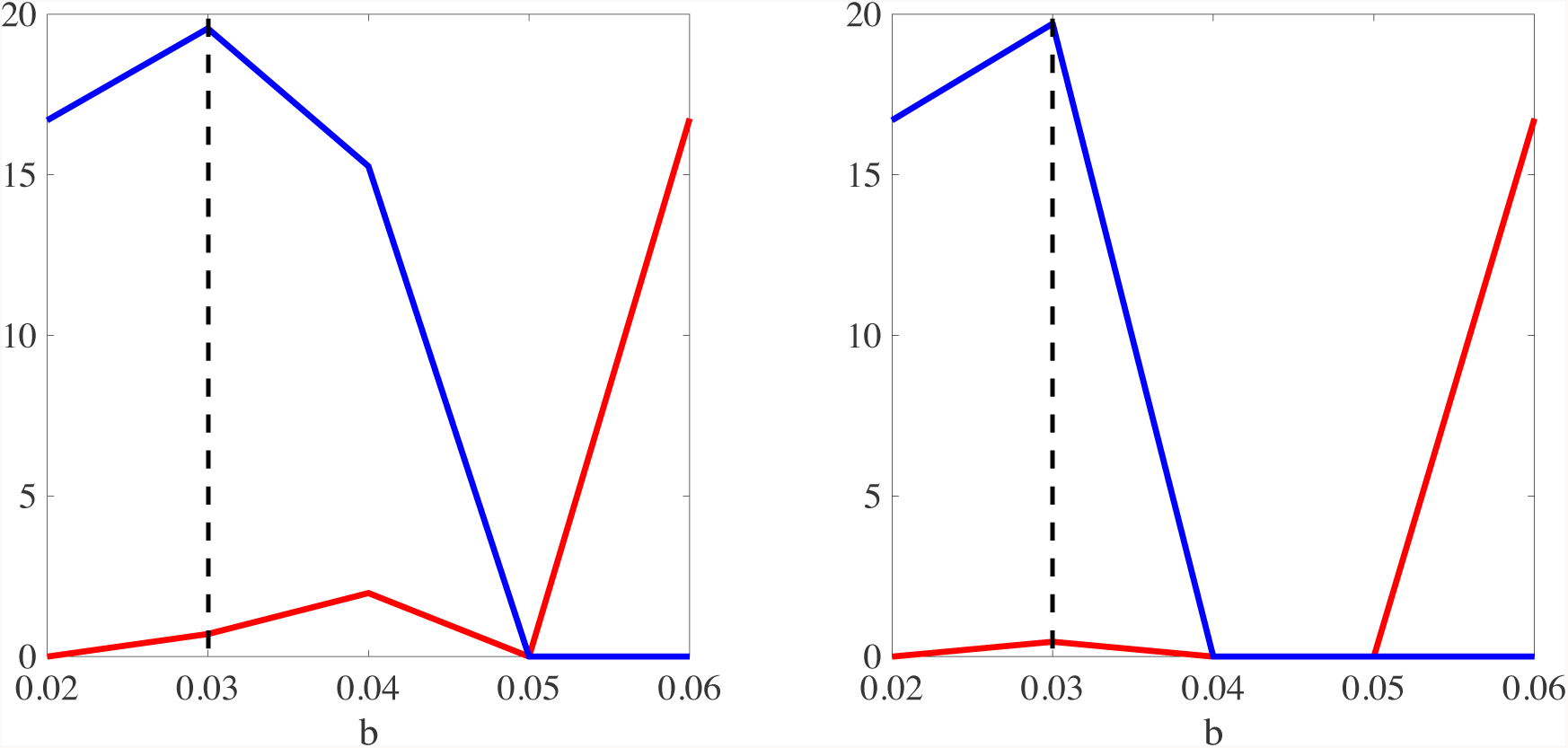
Plot of the time average of the total number of doves (blue lines) and hawks (red lines) as a function of the parameter *b*, in the absence (left panel) or in the presence (right panel) of semiochemical secretion and with non-synchronous generations. The black dashed lines highlight the value of the parameter *c* used to perform simulations. Time averages are in units of 10^4^.

### 3.3. Extension of the current modelling framework

In order to demonstrate the flexibility of the modelling framework presented here, which can be easily extended to span a broad spectrum of biologically relevant scenarios, we carry out computational simulations in the case in which the secretion rate of semiochemical can mutate over time. To do this, instead of modelling the rate of semiochemical secretion by means of the constant parameter *ν*, we associate to each player *i* a secretion rate 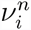 at the time step *n*. At the beginning of simulations, we set 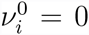 for all players (*i.e.,* for all values of *i*). We allow mutations to occur at reproduction and, focussing on the scenario in which mutations are small and rare, we let the size of mutations be either 0.01 with probability 0.001 or 0.1 with probability 0.0001. We run simulations for 2 × 10^6^ time steps. In the case of synchronous generations we assume *b* = 0.04 and *c* = 0.03, whilst we choose *b* = 0.03 and *c* = 0.04 in the case with non-synchronous generations. In both cases, the evolution of the total numbers of hawks and doves and the related time averages are actually very similar to those presented in the previous subsections for the case without semiochemical secretion, therefore we report here the plots of the total number of doves and hawks in the case of synchronous generations only in Figure 12 (these should be compared to the results in Figure 3). These results can be justified by noting that, at the end of simulations, the mean value of the secretion rate of semiochemical for doves and hawks are quite low, being 0.007 ± 0.04 and 0.143 ± 0.097 in the synchronous generation case, and 0.1415 ± 0.0901 and 0.136 ± 0.0729 in the non-synchronous generation case.

**Figure 12:**
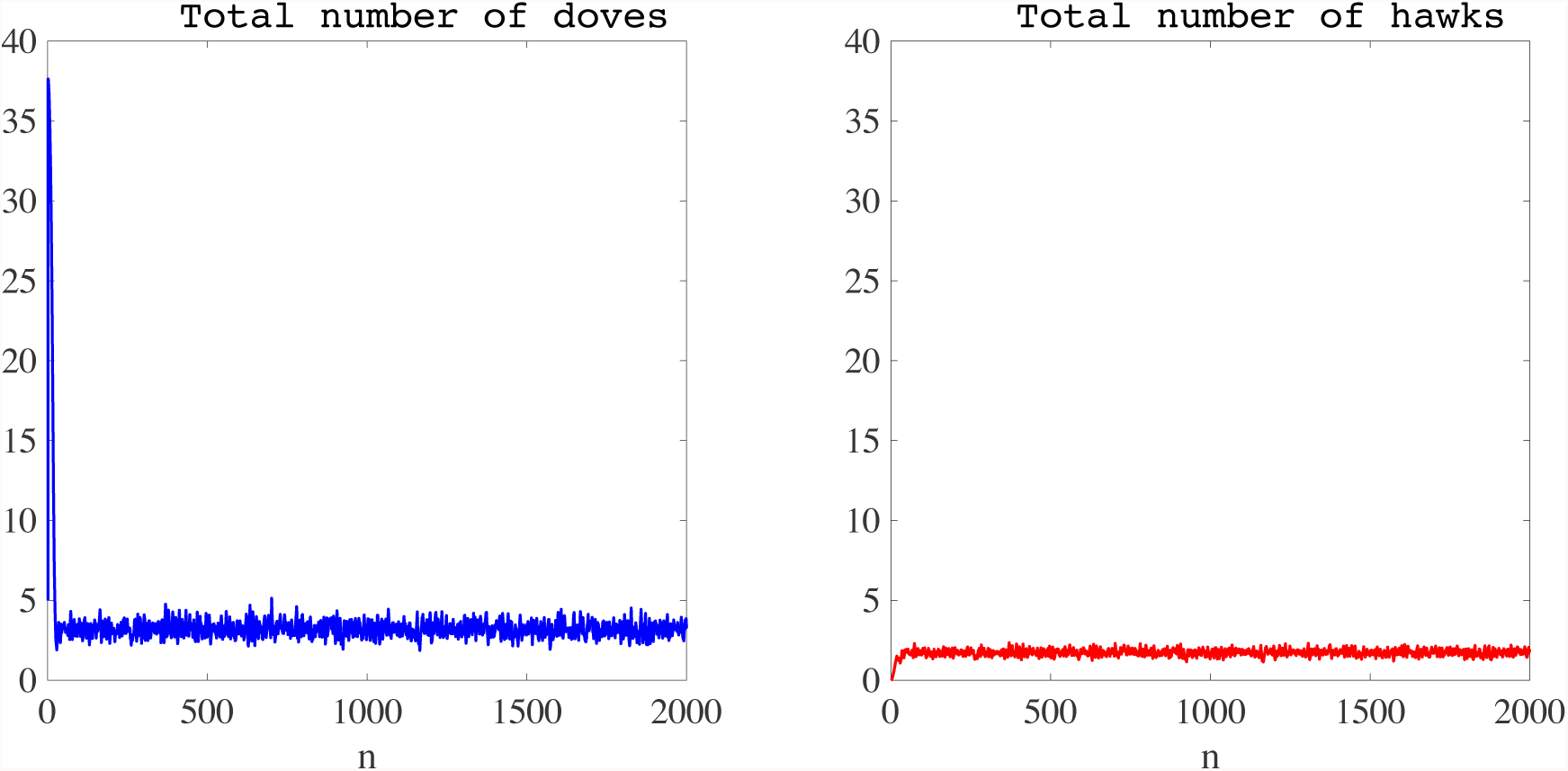
Plot of the total number of doves (left panel) and hawks (right panel) as a function of time, when mutations in the production rate of semiochemical occur and with synchronous generations. The time average of the total number of doves is 3.4288 × 10^4^, while that of hawks is 1.7193 × 10^4^. The time step *n* is in units of 10^3^, and the total numbers of doves and hawks are in units of 10^4^.

## 4. Discussion and conclusions

We have presented a hybrid modelling framework to study spatial evolutionary games. The originality of our approach stems from the fact that players explicitly move in space via random motion and chemotaxis.

In this framework, players occupying the same position can engage in a two-player game, and are awarded a payoff, in terms of reproductive fitness, according to their strategy. As an example, we have considered the Hawk-Dove game, for which it was previously shown that – both in the absence of any spatial structure [37] and when strategies move through colonisation of neighbouring sites [21] – if the benefit *b* exceeds the cost *c* then the hawk strategy is evolutionary stable, and doves will therefore be outcompeted by hawks. Our findings complement these previous studies by showing that, even if *b > c*, when one single hawk is introduced in a population of randomly scattered doves at time *t* = 0, allowing players to diffuse through space and perform chemotaxis can bring about self-organised dynamical patterns in which hawks and doves coexist in proportions that fluctuate about non-zero values. There is a striking difference between the patterns that emerge in the case where players’ generations are synchronous and the patterns observed in the presence of non-synchronous generations. In the former case, hawks become spontaneously organised into a travelling front which displaces the resident doves while expanding, whereas in the latter case hawks spread through the surrounding space leading to the formation of small holes (empty islands) in the distribution of doves. The waves of invasion created by hawks in the case of synchronous generations share some similarities with propagating waves arising in certain models of activator-inhibitor systems, excitable media and amoeboid aggregation, such as those reported by Erneux and Nicolis [14], Vasieva *et al.* [43] and Vasiev [42].

For synchronous generations, we have found that if *b* is sufficiently smaller than *c*, hawks are outcompeted by doves, which retain a competitive advantage because of their tendency to retreat at once if opponents escalate the conflict. On the other hand, for values of *b* sufficiently larger than *c*, doves are outcompeted by hawks. For intermediate values of *b*, we observe the stable coexistence of hawks and doves. The range of *b* values corresponding to stable coexistence is roughly the same both in the presence and in the absence of semiochemical secretion. Analogous considerations hold true for non-synchronous generations, although in this case there is a range of intermediate values of *b* for which coextinction of hawks and doves may take place. This range is wider in the presence of semiochemical secretion.

When coexistence occurs, we have shown that, compared with the situation without semiochemical, the presence of semiochemical sensing reduces the ratio between the time averages of the total number of hawks and doves, both for synchronous and non-synchronous generations. This result suggests that the chemotactic response to semiochemicals can be beneficial to doves, namely because it enhances the rate of conspecific interactions, and thus leads hawks to play a higher number of unproductive game rounds. Also, chemotactic response to the semiochemical alters the spatial patterns generated by doves and hawks in the case where generations are synchronous. In fact, when semiochemical is not present, doves form densely populated filamentary structures which are chased by relatively sparser filaments of hawks. Whereas, in the presence of semiochemical, doves become organised into dynamical clusters which are followed closely by flocks of hawks. On the other, when generations are non-synchronous the structure of patterns seems to remain essentially the same both in the presence and in the absence of semiochemical sensing.

We have made the assumption that the semiochemical is equally released from and sensed by all players, independently from their strategy. Also, we have assumed that: (i) the players cannot keep memory of past interactions; (ii) the offspring inherit the strategy of their parents, as no mutations occur; (iii) the repertoire of strategies available to players is limited to two. Such assumptions could be easily relaxed by virtue of the flexibility of our modelling framework, and the inclusion of these additional aspects can possibly produce a wider spectrum of spatial patterns. As a proof of concept, we have presented the results of numerical simulations carried out by allowing the players’ production rate of semiochemical to mutate over time.

While in this paper we have focused on the Hawk-Dove game, the modelling framework presented here can be used to explore the dynamics of any other spatial evolutionary game. For instance, we have already performed numerical simulations in the case where the players play either the Prisoner’s Dilemma game or the Rock-Paper-Scissors game. Overall, the results that we have obtained support the idea that our modelling framework allows for the study of unexplored scenarios in spatial evolutionary games. We also emphasise that we have not focussed on a specific system or bench-marked our model against any particular organisms. Rather the work presented here is intended to offer the perspective that mathematical modelling can complement more traditional methods of evolutionary biology research by capturing in qualitative terms the implicit assumptions of evolutionary hypotheses (and any hidden underlying assumptions) and clarifying the conditions under which certain evolutionary paths are possible. The generic modelling framework presented in this paper potentially covers a wide range of actual applications across a spectrum of biomedical systems including ecology (host-parasitoid systems, predator-prey systems; the colonisation of new habitats by invasive and, primarily, asexually reproducing plant species), epidemiology (the invasion of new host tissues by a pathogen, *e.g.*, the invasion of organs outside of the lungs by *Mycobacterium tuberculosis* [33]) and oncology (including the colonisation of a new niche by tumour cells following metastasis, the evolution of drug resistance [2, 3, 4]). Initial conditions that are far from the long-time limiting behaviour embody an organism encountering a new environment to which it is not currently phenotypically well-adapted, a rather ubiquitous scenario in biology and medicine.

In conclusion, we would like to remark that experience in a variety of contexts has demonstrated the value of relating individual-based stochastic models to deterministic continuum models [5, 7, 8, 10, 31, 32, 38], which make it possible to complement numerical simulations with rigorous analytical results, to achieve conclusions with broad structural stability under parameter changes. In this regard, a fundamental problem is what method of proof can be employed to derive deterministic mesoscopic models from the type of stochastic individual-based models which are defined in our framework. Addressing this fundamental question has potential to give unrivalled opportunities to link the phenotypic characteristics of single individuals with patterns of evolution and adaptation at the population level, and it can help to address pervasive biological problems concerning the mechanisms underlying the selection of behavioural traits in the presence of random motion and chemotaxis.

